# Predictive Models and Impact of Interfacial Contacts and Amino Acids on Protein-protein Binding Affinity

**DOI:** 10.1101/2023.09.07.555177

**Authors:** Carey Huang Yi, Mitchell Lee Taylor, Jesse Ziebarth, Yongmei Wang

**Author notes:** Corresponding author: Yongmei Wang, PhD.

## Abstract

Protein-protein interactions (PPIs) play a central role in nearly all cellular processes and that require proteins interact with sufficient binding affinity (BA) to form stable or transient complexes. Despite advancements in our understanding of protein-protein binding, much remains unknown about the interfacial region and its association with BA. Here we investigate the correlation of residue and atomic contacts of different types with BA and reveal the impact of the specific amino acids at the binding interface on BA. We create a series of linear regression (LR) models incorporating different contact features at both residue and atomic levels and examine how different methods of identifying and characterizing these properties impact the performance of these models. Particularly, we introduce a new and simple approach to predict BA based on the quantities of specific amino acids in contacts at the protein-protein interface. We show that the interfacial numbers of amino acids can be used to produce models with consistently good performance across different datasets, indicating the importance of the identities of interfacial amino acids in underlying the strength of BA. When trained on a diverse set of 141 complexes from two benchmark datasets, the best performing BA model (Pearson correlation coefficient R=0.68) was generated with an explicit linear equation involving six amino acids (tyrosine, glycine, serine, arginine, valine, and isoleucine). Tyrosine, in particular, was identified as the key amino acid in the quantitative link between specific amino acids and BA, as it had the strongest correlation with BA and was consistently identified as the most important amino acid in feature importance studies. Glycine, serine, and arginine were identified as the next three most important amino acids in predicting BA. The results from this study further our understanding of the importance of specific amino acids in PPIs and can be used to make improved predictions of BA, giving them implications for drug design and screening in the pharmaceutical industry.

## Introduction

Protein-protein interactions (PPIs) play a central role in nearly all cellular processes, from DNA replication to biomolecule transport and body immune response [1–4]. Abnormal protein-protein interactions are associated with many diseases such as cancer, neurodegenerative diseases, and infectious diseases [5–7]. Characterizing PPIs is therefore crucial for understanding biological processes and designing therapeutic drugs. PPIs are complicated processes, and it is the binding affinity (BA) of the interactions that determines whether proteins form stable or transient complexes and perform functions [8]. Although crystallography techniques are able to provide structural understanding of PPIs at the atomic level, these techniques do not directly measure BA.

BA is measured as the change in free energy (ΔG) during formation of a complex, where a negative value of a large magnitude indicates a strong interaction. ΔG is related to the dissociation constant (K_d_) by the formula ΔG = *R*T ln(K_d_) where *R* is the ideal gas constant, T is the temperature, and ΔG is the BA. BA can be measured in terms of dissociation constant (K_d_) using experimental approaches such as surface plasmon resonance (SPR) and isothermal titration calorimetry (ITC) [9]. In drug discovery, therapeutic design, and the development of inhibitors, predictions of the BA of proposed interactions are crucial for the identification of leading candidates [10]. For example, computational approaches, such as molecular docking, can reveal numerous theoretically possible protein-protein complexes within cells, and BA prediction is needed to evaluate which of these proposed structures are likely to be stable. However, the complexity of PPIs and the large diversity of protein structures and functions makes it difficult to predict BA and understand the features of the complex that impact BA.

During the past two decades, three general classes of computational approaches have been used to predict BA [11,12]: (1) physics-based ab-initio methods such as free energy perturbation and thermodynamics integration [13,14]; (2) knowledge-based statistical potentials such as DFIRE [15,16] or other empirical scoring functions such as Kdeep [17]; and (3) machine learning (ML)-based approaches based on structural and chemical properties of the protein-protein complexes [18–27] or their amino acid sequences [28–32]. Particularly, ML has become a sought-after approach for developing predictive models due to its high power of uncovering insights in large volumes of data [33]. The most common features in predictive ML models with 3D structures are the contacts at the complex interface, a method exemplified by Vangone and Bonvin with the development of the PRODIGY model and webserver in mid 2010s [19–21]. Compared to other ML-models, PRODIGY is simple and interpretable as it employs a linear regression (LR) algorithm, using the number of interfacial contacts (ICs) of polarity-classified residues and non-interacting surface (NIS) residue properties as the major features. The PRODIGY model achieved an excellent performance on a benchmark dataset of 81 complexes (Pearson’s correlation coefficient (R) of 0.73), outperforming numerous existing models. But, a recent study showed that PRODIGY’s performance on a different BA database was much lower (R = 0.31) [13], suggesting the need for further investigation of contact-based features and predictive models.

In this work, we compare a series of LR models involving residue/atomic contacts and then develop a new model based on quantitative information of the identities of the amino acids at the binding interface. As of today, there have been no studies that use the identities of specific amino acids to build models for BA prediction. Substantial studies have helped us to have a good understanding of the composition and role of specific amino acids at the complex interface for protein-protein binding [34–41]. However, translation of the information of the specific amino acids at binding interface into a BA prediction model has not been achieved. In this work, we report a quantitative association of the interfacial number (INs) of amino acids (AA) with BA and use this knowledge to develop a ML-based model for BA prediction. We found a group of high impact amino acids, led by tyrosine, that had an association with BA comparable to the most impactful residue or atomic contact features. We also show that this AA-INs model, which uses only six amino acids, outperforms other classes of contact-based LR models on benchmark datasets, indicating the importance of the identities of amino acids at protein interfaces in underlying the strength of protein-protein BA.

## Methods

### Features and their calculation

The contact features that we investigated included residue-based interfacial contacts (residue ICs) where a contact was defined between two residues if any of their heavy atoms were within a defined cutoff, atom-based interfacial contacts (atomic ICs) where any atoms that were within the cutoff were counted (not limited to one contact per residue), and hydrogen bonds (HB) where we used a cutoff of 3.5 Å between any N and O atoms. The residue ICs were further divided into different types based on charge and polarity of the amino acids. We used the following classifications: charged (Arg, Asp, Glu, His, Lys), polar (Asn, Gln, Ser, Thr), and apolar (Ala, Cys, Gly, Ile, Leu, Met, Phe, Pro, Trp, Tyr, Val). Atomic ICs were divided into three types based on the polarity of atoms, where C is considered as non-polar, and N, O, S as polar. We also considered the non-interacting surface (NIS) of the complex as it has been shown to contribute to BA [42]. NIS features were calculated as the percentage of surface area contributed by the polar, apolar, and charged residues over the total NIS area [19].

Additionally, we considered the hydrophobicity of the contacting residues by defining a hydrophobicity score (HS) for each complex. HS was defined according to:

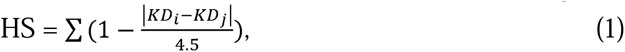

where the *KD* values for the two contacting residues, *i*th and *j*th, are the hydropathy indexes from the Kyte-Doolittle scale [43] and the score is summed over all pairs of contacting residues in the complex. The division by 4.5 and subtraction from 1 ensures that the score for each residue pair ranges between –1 and 1. With this definition, contacts between residues with similar hydrophobicity will contribute positive values near +1 to HS, while contacts between unlike residues have negative contributions to HS.

Thus, a total of 14 residue and atomic features were investigated and were indexed sequentially as: (0) residue ICs_charged-charged, (1) residue ICs_charged-polar, (2) residue ICs_charged-apolar, (3) residue ICs_polar-polar, (4) residue ICs_polar-apolar, (5) residue ICs_apolar-apolar, (6) HB, (7) atomic ICs_polar-polar, (8) atomic ICs_polar-apolar, (9) atomic ICs_apolar-apolar, (10) %NIS_polar, (11) %NIS_apolar, (12) %NIS_charged, and (13) HS. Residues ICs, atomic ICs, HB, and HS were calculated with Python, utilizing the Bio.PDB package within the Biopython library to extract the features from PDB files [44]. %NIS was calculated using NACESS with the same configurations used in the PRODIGY model by Vangone and Bonvin [19,45].

Additionally, the interfacial number (IN) of each amino acid (AA), AA-IN, for each complex was counted using the distance cutoff optimized by residue contacts. The AA-INs of the twenty standard amino acids were then used as features to develop LR models and investigate the impact of specific amino acids on BA.

The scripts for calculating these features and using the final AA-INs model based on six-amino acids for predicting BA of any protein complex with 3D structures are available at https://github.com/kagrat17/AAIN_Predictor. The code to calculate % NIS for the contact-based models was taken from the PRODIGY model by Vangone and Bonvin [20]. The distance cutoff for the residue and atomic contacts was optimized by investigating the Pearson’s correlation between the predicted ΔG and experimental ΔG at different distances.

### Datasets

To build structure-based models for predicting protein-protein BA, a diverse and reliable dataset of solved protein-protein structures with reliable affinity data is required. Vangone and Bonvin built a dataset by “cleaning” the structure-based protein-protein BA benchmark [46] and used it to develop the PRODIGY model and webserver [19,20]. This dataset, referred to as the PRODIGY dataset, contains 81 complexes with diverse functions and a wide range of experimental BA (ΔG from -18.6 to -4.3 kcal/mol). However, a later study by Romero-Molina et al. showed that PRODIGY did poorly on a different dataset that contained 90 complexes gathered from the PDBbind database (v.2020) [25,47]. PRODIGY gave a Pearsons’ correlation R of 0.31 on this PDBbind dataset in contrast to R of 0.73 on the PRODIGY dataset. In this work, we use both datasets as the resource of protein-protein complexes to develop improved models for BA prediction.

It has been shown by Vangone and Bonvin that the reliability of a dataset depends on the experimental methods used to determine K_d_ of the complexes [19]. The SPR, ITC, stopped-flow fluorimetry, and spectroscopic methods were shown to be reliable as the experimental ΔG measured by these methods gave reasonable correlations with residue ICs. Other methods, including inhibition assays and fluorescence spectrophotometry, gave BA with low correlations with residue ICs, possibly because they were indirect methods that were less reliable [48–50]. Complexes with K_d_ measured with these methods were thus removed by Vangone and Bonvin from the initial benchmark dataset. We conducted the same assessment for the PDBbind dataset and the results showed that the experimental ΔG from these other methods indeed gave low correlations with the number of residue ICs and atomic ICs (**Supplementary Table S1**). Thus, we excluded these complexes from the dataset. We also excluded two other complexes from the PDBbind dataset, 2FTL and 2JGZ. The experimental K_d_ for the complex 2FTL was measured at 100K rather than room temperature. The complex 2JGZ only had a lower bound K_d_ (K_d_ > 1mM). After removing these complexes, we had two final datasets of size 81 and 60, whose compositions are shown in **Supplementary Table S2**. We further examined whether there were significant differences between the two datasets. Specifically, we examined whether the BA distributions of their complexes were significantly different. The two-sample Kolmogorov-Smirnov test (testing whether two samples came from the same distribution) gave a p-value of 0.022, indicating that the distribution of BA differs between the two sets. This can also be seen by the BA histograms shown in **Supplementary Figure S1A**. Thus, we decided to combine the two datasets (total of 141 complexes) to make a large, diverse, and reliable dataset that can better represent protein-protein complexes. This combined dataset has a wide range of BA, with experimental ΔG from -3.3 to -18.6 kcal/mol (**Supplementary Figure S1B**).

### Machine learning methods

The statsmodels and sklearn python libraries were used to build and investigate BA models. We used different combinations of features to construct multivariate linear regressions and evaluated them using Pearson’s product-moment correlation coefficient R and the Akaike Information Criterion (AIC). In addition, root mean square error (RMSE) was calculated to evaluate the average difference between the predicted and experimental ΔG. We also built random forest (RF) models to rank the features of the models by their impurity-based feature importance. Cross-validation was also performed to evaluate our models by partitioning the dataset into four subsets, training on 75% of the data (training set) and validating on the other 25% of the data (validation set) and repeating until each of the 4 subsamples were used as the validation set. Such fourfold cross-validation was repeated 10 times, which gave mean test R with standard deviation.

## Results

### Sensitivity of contact-based prediction of BA on the distance cutoff used to define contacts

The simplest method of using protein-protein contacts to predict binding affinity is by creating a linear model using only the total number of interfacial contacts (ICs) of the residues from the two protein components. The total number of ICs increased rapidly with the distance cutoff used to define the contact (**Figure 1A**). To examine whether the model performance was sensitive to the cutoff used to calculate the number of ICs, we built LR models with the total number of ICs calculated at different cutoffs and examined the Pearson’s correlation R between predicted and experimental ΔG of these models. The value of R and, thus, the predictive ability of the number of ICs, increased as the cutoff distance increased above 3 Å, reaching a relative plateau from ∼4 to 6 Å, and then gradually decreasing as the cutoff continued to increase (**Figure 1B, black curve**). The maximum value of R occurred at a cutoff of 4.75 Å (R = 0.45), but the correlation was not particularly sensitive to the distance cutoff as distances of between ∼ 4 and 6 Å provided similar performance. Thus, a cutoff of 4.75 Å was used to calculate residue ICs for further studies.

**Figure 1.**
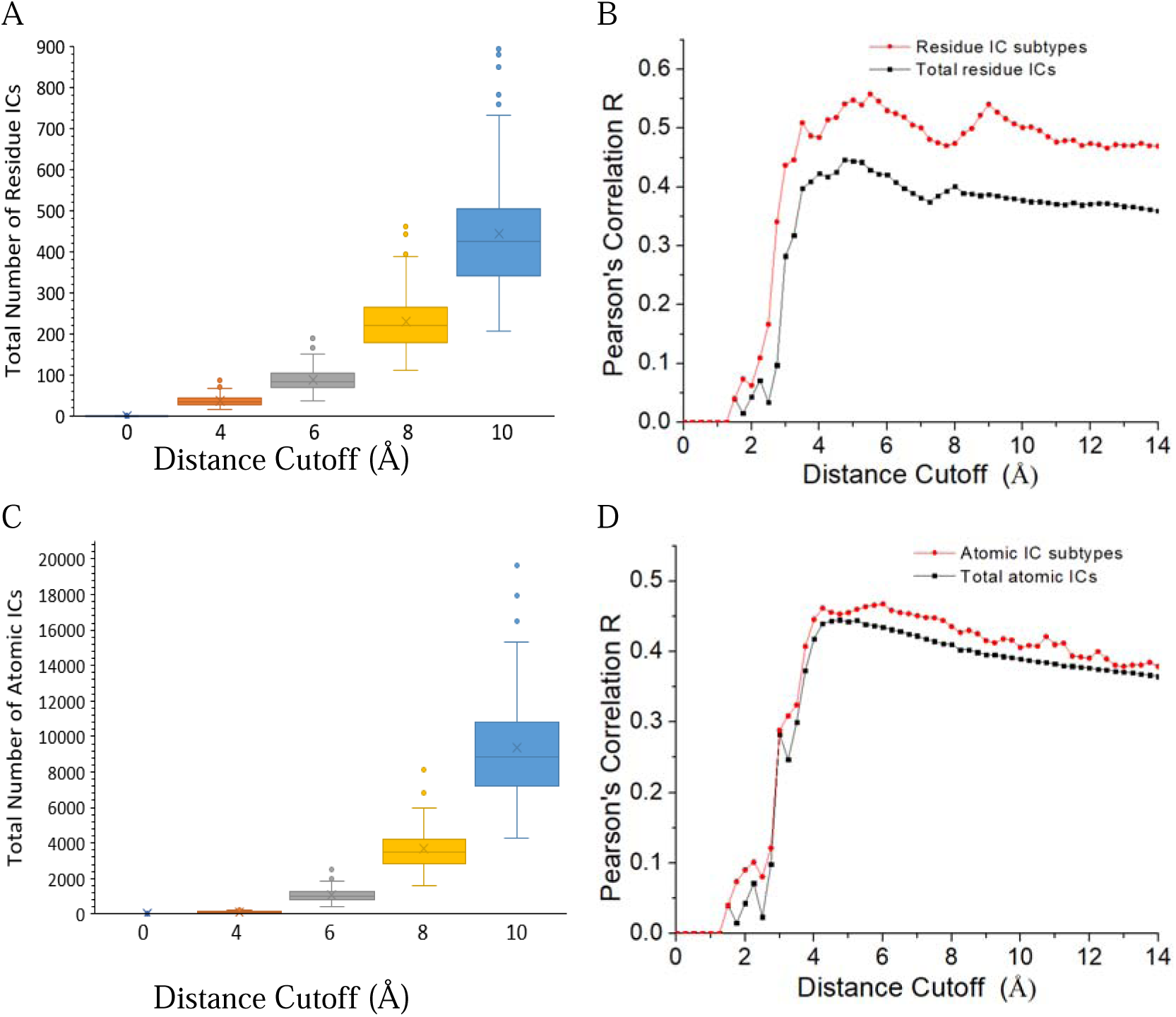
Dependence of the number of ICs and predictive performance of LR models on the distance cutoff and polarity-classified IC subtypes. (A) The total number of residue ICs at different distance cutoffs. (B) Comparison of the correlation between predicted and experimental ΔG at different distance cutoffs for the LR models generated with the features of total number of residue ICs (black curve) and the six residue IC subtypes (red curve). (C) The total number of atomic ICs at different distance cutoffs. (D) Comparison of the correlation between predicted and experimental ΔG at different distance cutoffs for the LR models generated with the features of total number of atomic ICs (black curve) and the three atomic IC subtypes (red curve).

Based on the polarity of residues, Vangone and Bonvin classified the residue ICs into six subtypes, ICs_charged-charged, ICs_charged-polar, ICs_charged-apolar, ICs_polar-polar, ICs_polar-apolar, and ICs_apopar-apolar, and showed that a LR model including subtype ICs performed better than the model using total ICs [19]. To examine whether this applies to a larger and more diverse dataset, we examined the relationship between the distance cutoff used to define a contact and the correlation between experimental ΔG and ΔG predicted by a LR model with these six features using the combined PRODIGY and PDBbind dataset. The general behavior of this relationship was similar to that found using only the total number of ICs, as R plateaued for cutoff distances between ∼ 4 to 6 Å and reached a maximum R of 0.54 at the same cutoff distance 4.75 Å (**Figure 1B, red curve**). However, in agreement with Vangone and Bonvin’s results, we found that the model performance improved when splitting the total contact number into the numbers of IC subtypes, with R values increasing by about 25%.

While the previous discussion has focused on contacts between the residues in protein-protein complexes, these residues actually interact with each other through atoms. Thus, we also investigated the correlation between the number of atom-atom contacts and binding affinity to compare residue-based and atomic-based approaches for defining protein-protein contacts. Defining contacts on an atomic basis greatly increased the number of contacts that were identified in the complexes, as there was over an order of magnitude increase in atomic contacts in the complexes compared with residue contacts (**Figure 1C**). The relationship between the cutoff distance used to define a contact and the correlation between the number of atomic contacts and the experimental ΔG (**Figure 1D, black curve**) was similar to that found for residu ICs, showing a plateau around 5 Å. The peak R for atomic ICs occurs at the same cutoff (4.75 Å) as that found for residue ICs, and the R value at the peak was similar for the two approaches (R = 0.44 for atomic ICs and R = 0.45 for residue ICs). Similar to how residue contacts can be classified based on the classes of the interacting residue, the atomic ICs can be grouped into three types (polar-polar, polar-apolar, and apolar-apolar) based on the polarity of atoms, where C is considered as non-polar, and N, O, S as polar. Different from the residue contacts, we found that the correlation of the LR models based on the three atomic subtypes was only slightly improved at each distance cutoff compared to those using the total atomic ICs (**Figure 1D, red curve**), with R only reaching 0.45 at the optimal distance cutoff, a value only slightly above that of the LR model with the total atomic ICs.

### Correlation between residue/atomic features and experimental BA

To further investigate and compare the relationship between residue/atomic ICs and BA, we calculated the Pearson’s correlation R between individual features and the experimental ΔG **(Table 1)**. The majority of these features had a negative correlation with ΔG and thus will lead to stronger BA. Several individual features were able to produce correlations with experimental ΔG comparable to total residue or total atomic ICs. Specifically, atomic ICs_polar-apolar, atomic ICs polar-polar, residue ICs polar-apolar, and HS all had R values of -0.4 or less, near the R ∼ -0.45 achieved by the total residue or atomic ICs. The strong correlation between experimental ΔG and residue ICs polar-apolar was previously reported by Vangone and Bonvin with the PRODIGY dataset [19]. Other features with relatively strong correlations with experimental ΔG included residue ICs charged-apolar, atomic ICs apolar-apolar, %NIS_polar, and HB. %NIS_charged had a relatively strong positive correlation with ΔG (R = 0.30), indicating that increasing the percentage of NIS from charged residues would lead to weaker BA.

**Table 1.** Pearson’s correlation R between the experimental ΔG and residue ICs, atomic ICs, HB, HS, and %NIS features.

| **Residue-ICs** | **R** | ***p-value*** |
| --- | --- | --- |
| Residue ICs_charged-charged | -0.12 | 0.078 |
| Residue ICs_charged-polar | -0.21 | 0.006 |
| Residue ICs_charged-apolar | -0.39 | <0.0001 |
| Residue ICs_polar-polar | -0.15 | 0.038 |
| Residue ICs_polar-apolar | -0.44 | <0.0001 |
| Residue ICs_apolar-apolar | -0.19 | 0.012 |
| Total residue ICs | -0.45 | <0.0001 |
| HS | -0.41 | <0.0001 |
| **Atomic ICs** |  |  |
| Atomic ICs_polar-polar | -0.44 | <0.0001 |
| Atomic ICs_polar-apolar | -0.45 | <0.0001 |
| Atomic ICs_apolar-apolar | -0.37 | <0.0001 |
| Total atomic ICs | -0.45 | <0.0001 |
| HB | -0.33 | <0.0001 |
| **%NIS** |  |  |
| %NIS_polar | -0.35 | <0.0001 |
| %NIS_apolar | 0.09 | 0.144 |
| %NIS_charged | 0.30 | 0.0002 |

One somewhat surprising result from this analysis is that contacts between ‘like’ residues or ‘like’ atoms (e.g., polar-polar residue contacts or apolar-apolar atomic contacts) had weaker correlations with BA than ‘unlike’ contacts. For example, polar-apolar residue and atomic contacts had the strongest correlation with experimental ΔG in their respective categories, indicating that these ‘unlike’ interactions were the best at predicting BA. To further investigate ‘like’ and ‘unlike’ contacts, we calculated the expected percentage (average from 141 complexes) of residue pairs in contacts based on the percentages each residue class in contacts and compared these expected percentages with their actual values (**Supplementary Table S3**). All ‘like’ residue pairs (charged-charged, polar-polar, and apolar-apolar) had higher percentages than their expected values by ∼10%. Thus, there does seem to be a preference for the formation of ‘like’ contact pairs at the protein interface, but this preference does not translate to these contacts being better predictors of binding affinity.

### Comparison of residue, atomic, and combined feature sets in creating LR models of BA

To more fully explore the contact-based features and predictive models, we built and evaluated a series of LR models with different feature combinations. First, we used the set of features used in the development of the PRODIGY model to build residue ICs/NIS models to examine how these models performed on the expanded dataset used in this work that contained the 81 complexes from the PRODIGY dataset and additional 60 complexes from PDBbindt. Specifically, the features used here included six types of residue ICs (Feature Index 0,1,2,3,4,5) and three %NIS (Feature Index 10,11,12). We trained a total of 511 LR models using all possible combinations of these nine features on our dataset. The performance of these models was evaluated by both Pearson’s correlation R (**Figure 2A**) and AIC (**Figure 2B**). The AIC criteria was used to correct for overfitting and gave more weight to simpler models. The results showed that increasing the number of the features from 1 to 3 led to a rapid increase of the model performance. The maximal *R* (*R*_max_) increased by 0.15 and the minimal AIC (AIC_min_) decreased by 26.1 from single-feature models to three-feature models. However, as the number of features included in the model increased above 5, the *R*_max_ had negligible changes (remaining near 0.61), while the AIC_min_ increased as AIC penalized the increasing complexity of these models. This residue ICs/NIS model with the lowest AIC (AIC = 622.0) gave R of 0.61 (p < 0.0001) and RMSE of 2.10 kcal/mol (**Figure 2C**). It included the following 5 features: residue ICs_charged-charged, residue ICs_charged-apolar, residue ICs_polar-polar, residue ICs_polar-apolar, and %NIS_polar. While this model had the lowest AIC, many models performed relatively well, with 153 of the 511 models giving R > 0.55.

**Figure 2.**
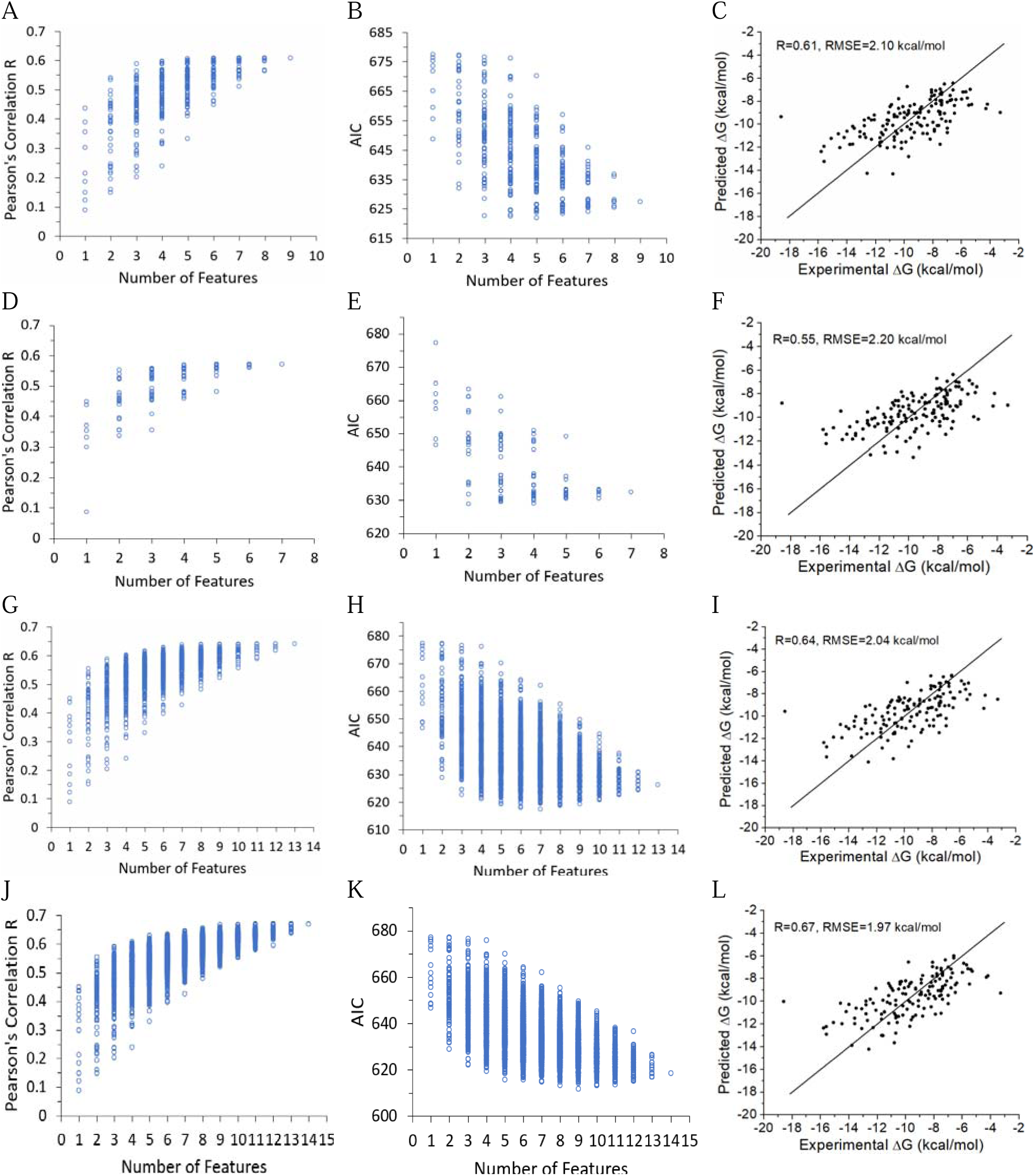
Comparison of the performance of different groups of contact-based LR models. (A-C) Residue ICs/NIS models. (D-F) Atomic ICs/NIS models. (G-I) Residue/atomic ICs/NIS models. (J-L) Residue/atomic ICs/HS/NIS models. (A, D, G, J) Pearson’s correlation R as a function of the number of features used to generate the models. (B, E, H, K) AIC as a function of the number of features used to generate the models. (C, F, I, L) Scatter plots between the predicted and experimental ΔG for the minimum AIC models. The straight lines are the function *y=x*.

Next, we built a set of LR models based on atomic contacts to compare with residue-based models. These atomic contact-based models included all possible combinations of 7 features, HB, atomic ICs_polar-polar, atomic ICs_polar-apolar, and atomic ICs_apolar-apolar features, and the three %NIS features (Feature Index 6,7,8,9,10,11, 12), resulting in a total of 127 LR models. The changes of R and AIC as a function of feature number for these atomic IC/NIS models had similar trends as residue ICs/NIS models, as R_max_ and AIC values soon reached a plateau as more than two features were added to models (**Figure 2D** and **Figure 2E**). The R_max_ increased from 0.45 for single-feature model to 0.55 for two-feature model, and only increased to 0.57 for the all-feature model. The model with the minimum AIC (AIC=628.8) included two features, ICs_polar-apolar and %NIS_polar and gave R = 0.55 (p < 0.0001) and RMSE = 2.20 kcal/mol (**Figure 2F**). Again, many atomic ICs/NIS models (>20%) performed similarly to the minimum AIC model.

Then, we examined whether combining residue-based and atomic-based contact features of protein complexes would improve predictions of their experimental ΔG. The combination of residue contacts, atomic contacts, and %NIS features resulted in a total of 13 features and 8191 possible models. Both R and AIC improved significantly as more features were added until 7 features, with R_max_ remaining at 0.64 for models with 7 to 13 features **(Figure 2G** and **Figure 2H)**. The model with the lowest AIC (AIC = 617.3) included a total of 7 features and gave R = 0.64 (p < 0.0001) and RMSE = 2.04 kcal/mol (**Figure 2I)**. The 7 features in this model included a mix of atomic ICs, residue ICs, and %NIS, which were residue ICs_charged-polar, residue ICs_charged-apolar, residue ICs_polar-polar, residue ICs_polar-apolar, HB, atomic ICs_polar- polar, and %NIS_polar. Similarly, many residue/atomic ICs/NIS performed relatively well, with ∼ 20% of the models giving R > 0.60.

One additional way to characterize residue interactions is to use hydropathy indices of residues. Hydropathy index is a number representing the hydrophobic or hydrophilic properties of its side chain. The larger the number is, the more hydrophobic the amino acid is. Several hydrophobicity scales have been published. We used the commonly used Kyte-Doolittle scale [43] to calculate the hydrophobicity score, HS, for each protein-protein complex. Adding HS to the residue ICs, atomic ICs, and %NIS gives a total of 14 features, resulting in 16,383 possible feature combinations to create models. R reached the maximum of 0.67 for models with 9 to 14 features, and the AIC reached minimum of 611.7 at 9 features (**Figure 2J** and **Figure 2K**). About 30% of the residue/atomic ICs/HS/NIS models gave R > 0.60. The minimum AIC model used 9 features: residue ICs_charged-polar, residue ICs_charged-apolar, residue ICs_polar-polar, residue ICs_polar-apolar, residue ICs_apolar-apolar, HB, atomic ICs_polar-polar, % NIS_polar, and HS, with R of 0.67 (p < 0.0001) and RMSE of 1.97 kcal/mol (**Figure 2L**).

The minimum AIC models for the four different groups of features were tested by fourfold cross-validation and compared with the all-feature model for each class (**Table 2**). In all cases, the minimum AIC models performed slightly better than the all-feature models during cross-validation. For example, the test R of the minimum AIC residue/atomic ICs/HS/NIS model was 0.61 whereas the test R of the all-feature model was 0.58. The four different feature groups (i.e., residue ICs/NIS, atomic ICs/NIS, residue/atomic ICs/NIS, and residue/atomic ICs/HS/NIS) provided roughly similar results, as the R values ranged between 0.55 and 0.61. Thus, increasing from the two features included in the atomic ICs/NIS model to the nine-feature atomic/residue ICs/HS/NIS model only increased R from 0.54 to 0.61. The coefficients for the linear equations of the minimum AIC models and the all-feature models with their p-values are shown in **Table 3** and **Supplementary Table S4**, respectively. Generally, the minimum AIC model picked up features that had *p* < 0.05 in the all-feature models.

**Table 2.**
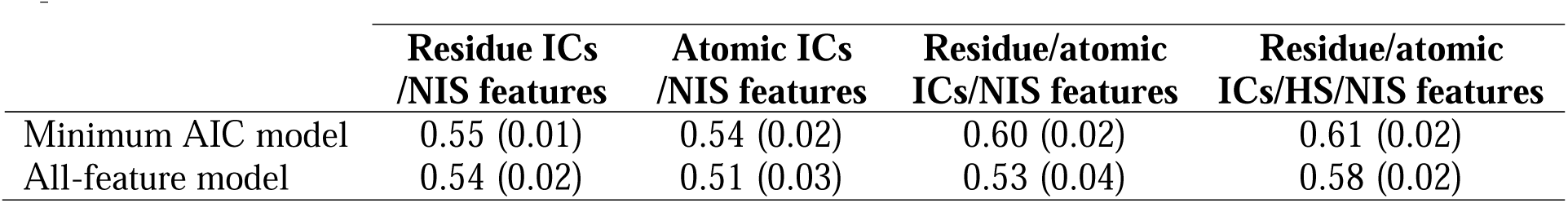
Test R for the cross-validation of minimum AIC and all-feature models. Data are presented as the mean with standard deviation from 10x trials.

**Table 3.** Coefficients of features in the linear equations predicting ΔG for minimum AIC models.

| Feature | Residue ICs<br>/NIS model |  | Atomic ICs<br>/NIS model |  | Residue/atomic ICs<br>/NIS model |  | Residue/atomic ICs<br>/HS/NIS model |  |
| --- | --- | --- | --- | --- | --- | --- | --- | --- |
|  | Coefficient | p-value | Coefficient | p-value | Coefficient | p-value | Coefficient | p-value |
| Residue ICs_charged-charged | -0.0735 | $p=0.107$ | | | | | | |
| Residue ICs_charged-polar | | | | | 0.1106 | $p=0.092$ | 0.1564 | $p=0.019$ |
| Residue ICs_charged-apolar | -0.1262 | $p=0.002$ | | | -0.1024 | $p=0.014$ | -0.1240 | $p=0.004$ |
| Residue ICs_polar-polar | 0.1456 | $p=0.154$ | | | 0.1811 | $p=0.042$ | 0.2689 | $p=0.004$ |
| Residue ICs_polar-apolar | -0.1528 | $p=0.001$ | | | -0.0947 | $p=0.032$ | -0.1171 | $p=0.013$ |
| Residue ICs_apolar-apolar | | | | | | | 0.0601 | $p=0.071$ |
| HB | | | | | 0.0893 | $p=0.117$ | 0.1232 | $p=0.031$ |
| Atomic ICs_polar-polar | | | | | -0.0680 | $p=0.003$ | -0.0588 | $p=0.010$ |
| Atomic ICs_polar-apolar | | | -0.0181 | $p=0.000$ | | | | |
| Atomic ICs_apolar-apolar |  |  |  |  |  |  |  |  |
| % NIS_polar | -0.1329 | $p=0.000$ | -0.1242 | $p=0.000$ | -0.1241 | $p=0.000$ | -0.1198 | $p=0.000$ |
| % NIS_apolar |  |  |  |  |  |  |  |  |
| % NIS_charged |  |  |  |  |  |  |  |  |
| HS | | | | | | | -0.1102 | $p=0.004$ |
| Constant | -1.796 | $p=0.000$ | -2.393 | $p=0.000$ | -1.925 | $p=0.000$ | -2.131 | $p=0.000$ |

### Feature importance studies revealed the impact of different residue/atomic properties on BA

To decipher interfacial factors that impact BA, we conducted feature importance studies with several approaches. First, we examined the feature composition in the minimum AIC models for different feature combinations (**Supplementary Figure S2**). We then calculated the frequency of feature occurrence in the top models (**Figures 3A - 3D**). We chose a minimal number of the top models such that each feature gave a different frequency of occurrence in these models. This was top 105 for the residue ICs/NIS models, top 50 for the atomic ICs/NIS models, top 100 for the residue/atomic ICs/NIS models, and top 200 for the residue/atomic ICs/HS/NIS models. In addition, we predicted the feature importance by RF using the all 14- feature model (**Figure 3E**). For convenience, the feature index was used in all the plots.

**Figure 3.**
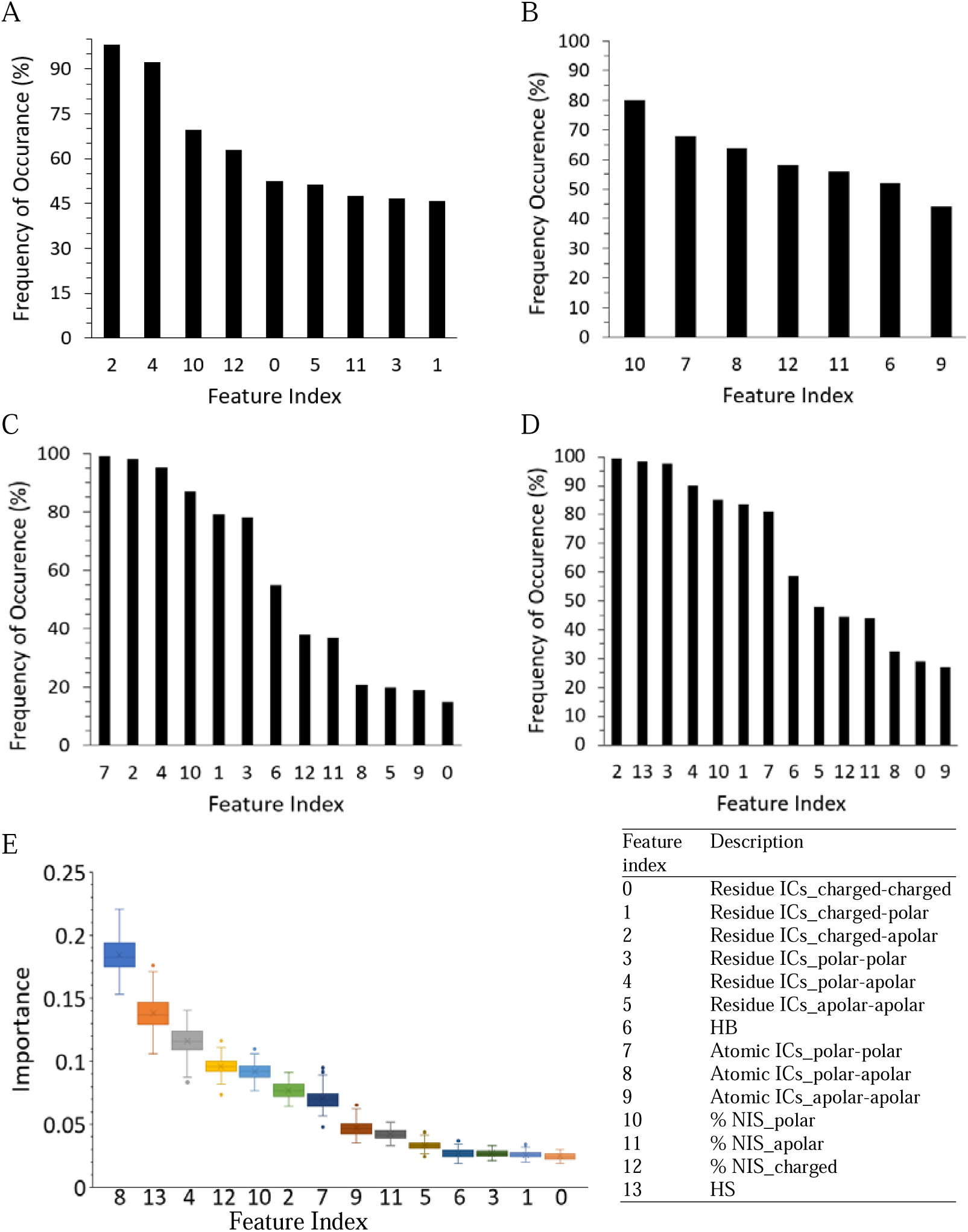
Feature importance of residue ICs, atomic ICs, HB, HS, and %NIS by frequency of occurrence in top models (A-D) and RF (E). (A) Residue ICs/NIS models. (B) Atomic ICs/NIS models. (C) Residue/atomic ICs/NIS models. (D) Residue/atomic ICs/HS/NIS models. (E) Feature importance predicted by RF using the all-feature model.

Due to the different features used and the different methods used to identify important features, there were, not surprisingly, large differences in the features that were identified as most important for each feature set. Additionally, there are high correlations between many of the contact features (**Supplementary Table S5**). For example, the atomic ICs_polar-polar had high correlation with atomic ICs_polar-apolar (R = 0.93), residue ICs_charged-polar (R = 0.74) and hydrogen bond (R = 0.90). Therefore, the addition of these three features to a model that contained atomic ICs polar-polar would involve redundant information and would not be likely to significantly improve the model. However, a couple of consistent results were identified. First, HS was identified as the second most important feature in both cases when it was included in the model, indicating that including the hydropathy index of contacting residues contained valuable information for predicting experimental ΔG. On the other hand, HB was not identified as one of the five most important features in any of the models and was always ranked behind atomic polar-polar contacts in importance. While HB was included in the minimum AIC model for two feature sets (**Table 3**), it had a lower p-value than atomic polar-polar contacts in both models. Finally, in agreement with our previous discussion on the R values for single feature LR models, feature importance between ‘like’ pairs of residues and contacts did not seem to consistently be more important features than ‘unlike’ contacts. For example, residue contacts between apolar and either charged or polar residues (Feature Index 2 and 4) were usually identified as being the most important class of residue contacts.

### Interfacial numbers of amino acids, enrichment and depletion

Inspired by our analysis of the importance of contact features (e.g., the high importance of polar-apolar residue and atomic contacts) and recent results that have shown the importance of specific residues such as tyrosine at the binding interface in protein-protein complexes [36], we further investigated the amino acids that were found at the binding interface. Specifically, we determined the enrichment or depletion of amino acids at the binding interface compared to their presence in the entire complex (**Figure 4** *and* **Supplementary Table S6**).

**Figure 4.**
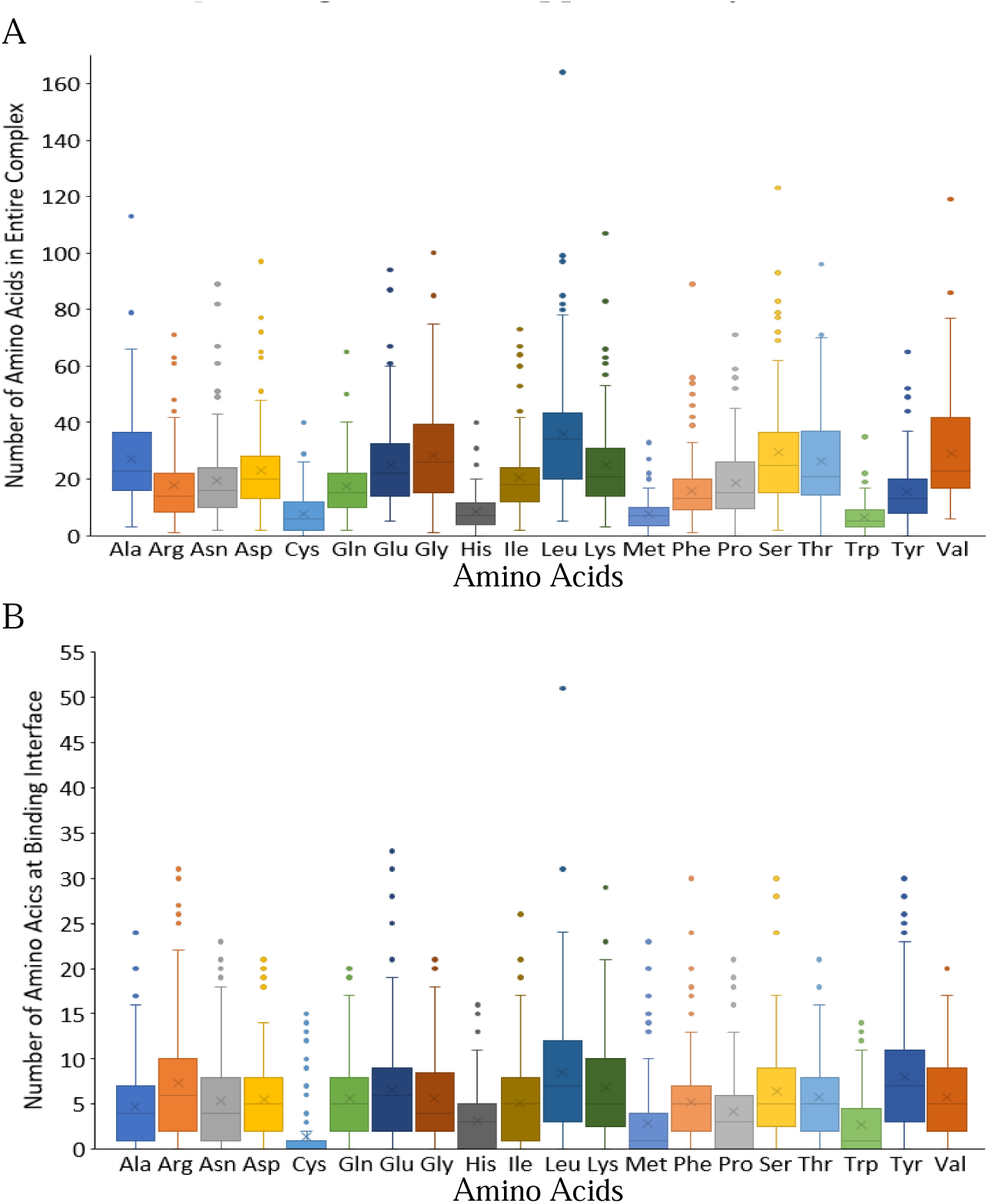
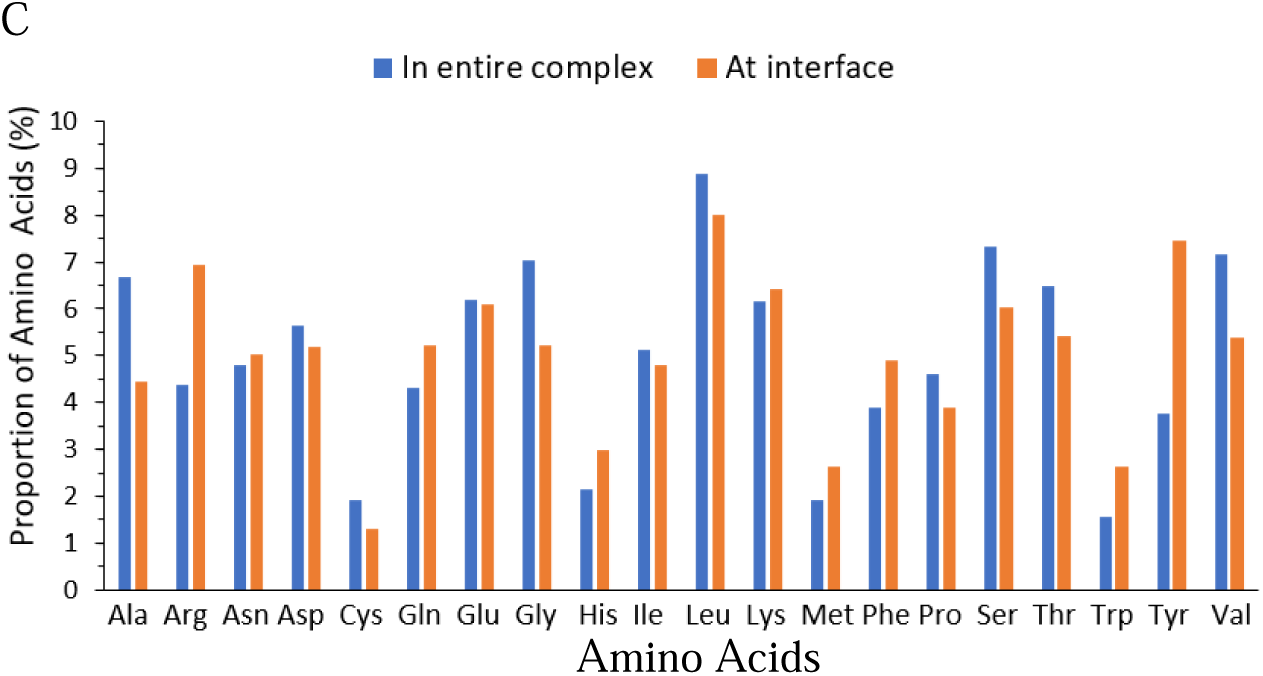
Abundance of the amino acids in the entire protein-protein complex and at the binding interface. (A) Box plots of the number of each amino acid in the entire protein-protein complex in the dataset. (B) Box plots of the number of each amino acid involved at the binding interface (AA-IN) of protein-protein complex in the dataset. (C) Comparison of the proportion of amino acids at the interface and in the entire complex.

First, we calculated the number and proportion of each amino acid in the entire complex. Next, we calculated the number of each amino acid at the binding interface, AA-IN and their proportions at the optimal cutoff of 4.75 Å. An enrichment factor was then calculated using the ratio of these two percentages (% at interface/% in entire complex), where enrichment factors > 1 indicate that an amino acid had an abundance at interface that was greater than expected by its abundance in the entire complex.

The results showed that leucine was the most abundant amino acid in protein-protein complexes, accounting for 8.9% of all amino acids. The least abundant amino acid was tryptophan, which only accounted for 1.6%. These results agree with those by others using large datasets (Swissprot and TrEMBL) that showed leucine was the most abundant amino acid whereas tryptophan and cystine are the least abundant [39], indicating that the amino acid percentages in these protein-protein complexes in our dataset are similar to those found in all proteins.

Of these residues at the binding interface, leucine remained the most abundant amino acids (8.0%), with Leu-IN reaching as high as 51 for the 5YR0 complex. However, the enrichment factor of leucine was < 1, as it was more prevalent in entire complex than it was in the contacts. Tyrosine was the second most abundant amino acid at the binding interface, accounting for 7.5% of all amino acids at the interface. Compared to its abundance in the entire complex (3.8%), the abundance of tyrosine at the interface was doubled, making it the amino acid with the highest enrichment factor. For individual complex, the highest percentage of tyrosine at interface was 28.3% (2AJF). Other remarkably enriched amino acids at the contact interface were arginine and tryptophan which had enrichment factors near 1.65. Alanine had th lowest enrichment factor among the twenty amino acids. It was the fifth most abundant amino acid in the entire complex, but it became the 15th at contact interface.

### AA-INs are correlated with experimental BA

To examine the association of specific amino acids with BA, we determined whether the numbers of individual amino acids at the interface (AA-INs) in each complex were correlated with the experimental ΔG of the complex (**Table 4**). The R values of the correlations between ΔG and the INs of several amino acids were comparable to the best performing residue and atomic ICs features. Specifically, the INs of tyrosine had a lower R value (R = -0.49, p < 0.0001) than any of the previously discussed contact features, while glycine (R = -0.45, p < 0.0001) and serine (R = -0.44, p < 0.0001) were able to perform as well as any of the residue and atomic ICs features when predicting experimental ΔG. Tyrosine has been shown to play a dominant role for protein-protein binding due to its unique physicochemical properties that make it effective at mediating molecular recognition [36]. Thus, simply using the number of tyrosine residues involved in contacts at the interface of a protein-protein complex is better able to predict the BA of the complex than other features including the total number of atomic or residue ICs or the number of any one of the IC contact subclasses. We also investigated the correlation between ΔG and enrichment factor (**Table S7**). However, these correlations were lower than those of AA- INs and thus we decided to focus our investigation on AA-INs.

**Table 4.**
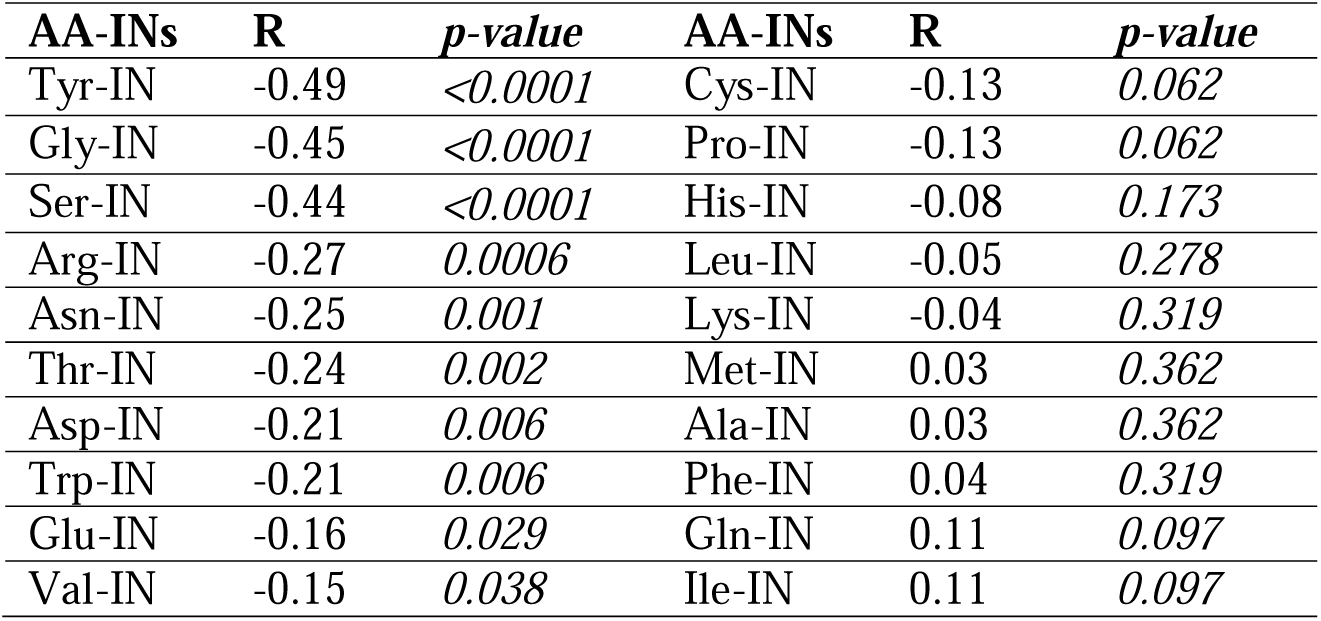
Pearson’s correlation R between AA-INs and experimental ΔG.

### AA-INs based models predict BA with accuracy comparable to contact based models

To investigate the feasibility of specific amino acids for BA prediction, we used the numbers of each amino acid at the binding interface, AA-INs, of the twenty amino acids to build LR models predicting BA and understand the relationship between specific amino acids and BA. Using the AA-INs of the twenty amino acids, we built 1,048,575 LR models for all possible combinations of 20 features. Due to the huge number of models, we selected the top 2000 models to examine the R and AIC as a function of feature combination. These top 2000 models included models with the number of features between 4 and 13 (**Figure 5A** and **Figure 5B**). Similar to the results for the residue and atomic IC models above, the R_max_ increased as the number of features were added, but AIC reached a minimum of 600.9 when six features were used. This minimum AIC model used the following six amino acids: tyrosine, glycine, serine, arginine, valine, and isoleucine with a simple linear equation of

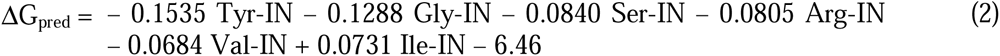

**Figure 5.**
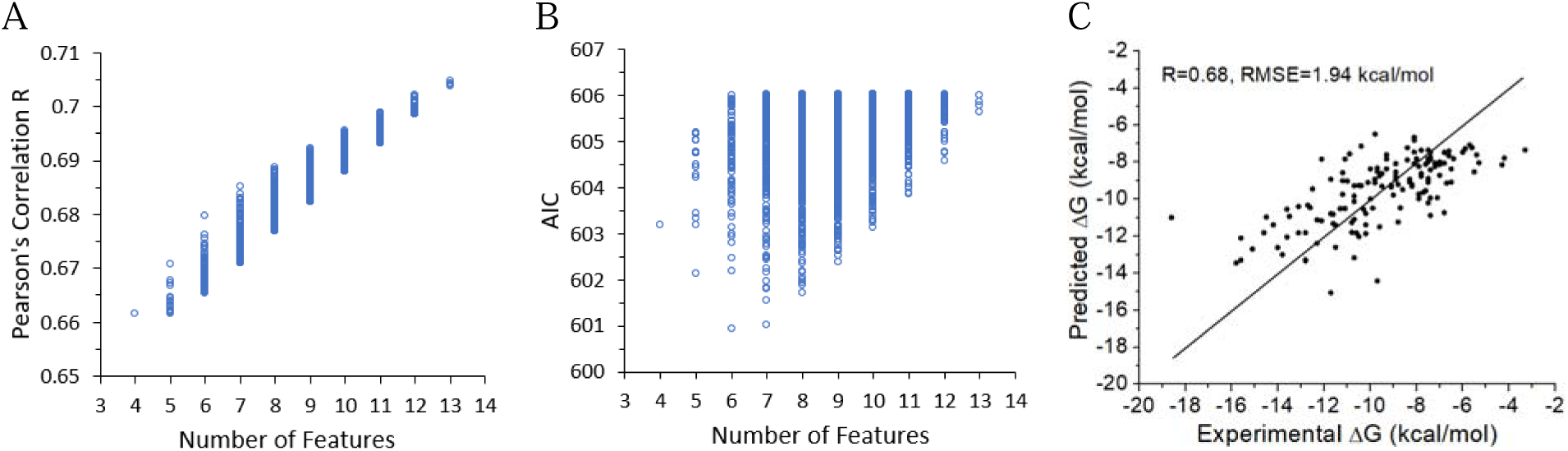
Performance of the AA-INs models for BA prediction. (A) Pearson’s correlation R and (B) AIC as a function of the number of features used to generate the top 2000 models. (C) Scatter plot for the minimum AIC AA-INs model between predicted and experimental ΔG. The straight line is the function *y=x*.

This model gave R = 0.68 (p < 0.0001) and RMSE = 1.94 kcal/mL, with test R = 0.63 in cross- validation (**Figure 5C** and **Supplementary Table S8**). This model outperformed all of the models above that used a combination or a subset of residue ICs, atomic ICs, HS, and NIS features, while using only six features, as compared to the nine features used in the residue/atomic ICs/HS/NIS model. To examine whether the model can be further improved by adding residue and atomic ICs and the high impacting NIS and HS features, we made sets of models combining AA-INs with these features. The best performing model gave an R of 0.71 and a test R of 0.67 after cross-validation, indicating that combining AA-INs with these other features does not greatly improve predictions.

The six amino acids included in the minimum AIC AA-INs model included four apolar (tyrosine, glycine, valine, isoleucine), one polar (serine), and one charged (arginine) residue. Thus, differences in the relative importance of the different amino acids may help explain the lack of agreement between residue similarity and feature importance that we observed when analyzing LR models based on residue ICs. For example, a residue IC that contained an ‘unlike’ combination of residues such as tyrosine and threonine would be more likely to have a strong impact on binding affinity than a ‘like’ combination of residues with INs that are not correlated with BA. To support this conclusion, we investigated the number of times that tyrosine was in contact with each of the other amino acids and found that, of the 5 most common tyrosine contact partners, 2 were polar (asparagine and threonine) and 2 were charged (lysine and arginine). These ‘unlike’ contacts that included tyrosine would increase the relative importance of polar-apolar and charged-apolar ICs, making these features more important in general when analyzing the residue contact - based LR models.

### Feature importance studies identified a group of amino acids with high impact on BA

To investigate the importance of the twenty amino acids in BA prediction, we examined how the features were used to build the top 1000 models via the frequency of feature occurrence in these models. The results showed that there were six amino acids that were used with much higher frequency than others in the top 1000 models, tyrosine, glycine, arginine, serine, isoleucine, and valine (**Figure 6A)**. These top six amino acids were the same six amino acids that were used to generate the minimum AIC model. Particularly, tyrosine, glycine, and arginine were used in all the top 1000 models.

**Figure 6.**
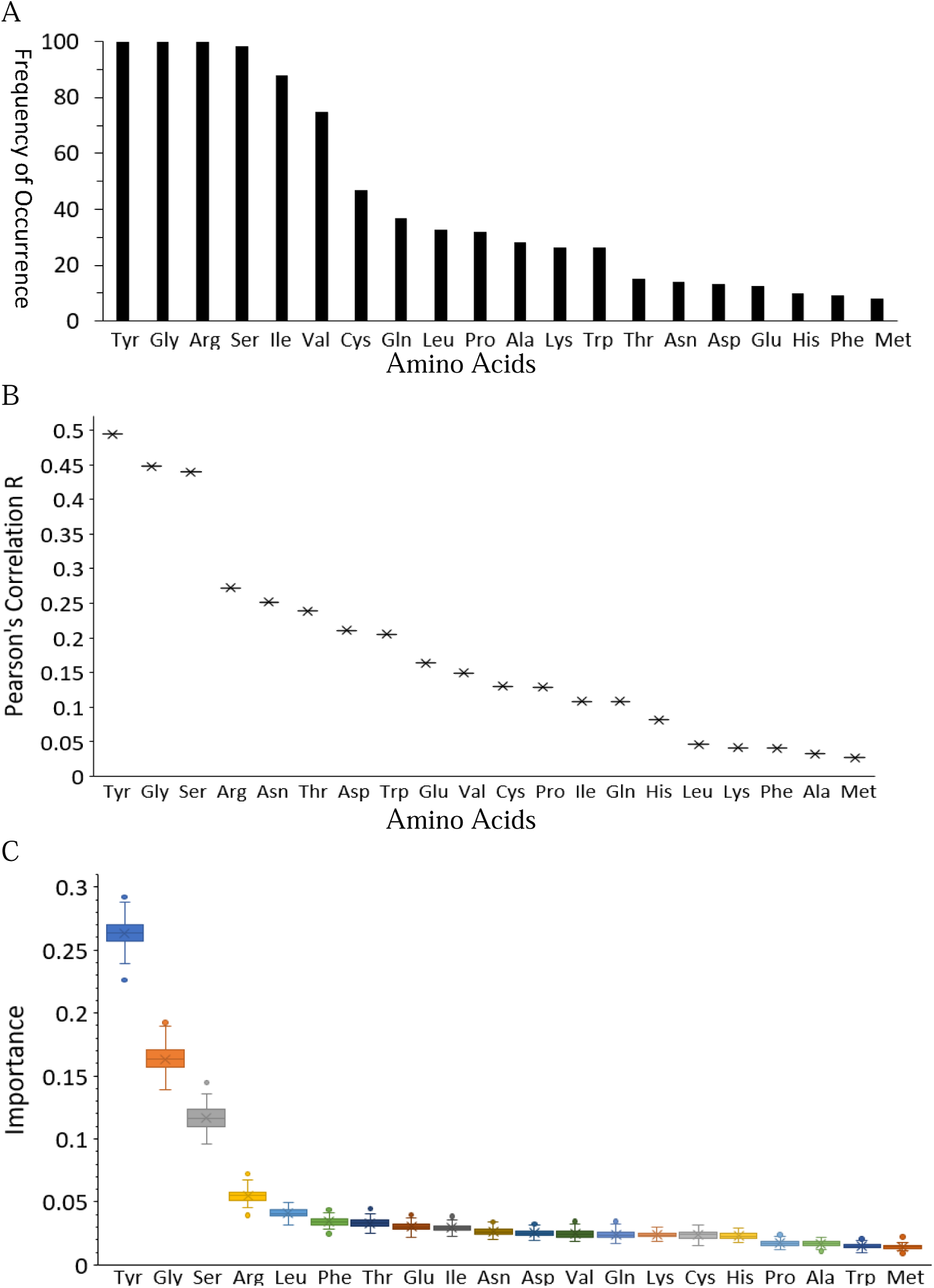
Feature importance of AA-INs by different approaches. (A) Frequency of occurrence in the top 1000 AA-INs models. (B&C) Feature importance of the twenty amino acids predicted by one-parameter LR (B) and RF (C).

Additionally, feature importance was studied with one-parameter LR modeling and RF. The one-parameter LR ranked tyrosine as the leading amino acid (R = 0.49), followed by glycine (R = 0.45) and serine (R = 0.44) (**Figure 6B**). These three amino acids performed much better than the rest of amino acids (R < 0.3) in BA prediction. Arginine was ranked the fourth important feature with R = 0.27. However, the other two top amino acids given by their frequency of occurrence in the top 1000 models, isoleucine and valine only gave R of 0.11 and 0.15 respectively. The amino acid that gave lowest R was methionine, with R of only 0.027. Methionine was also the least frequently used amino acid in the top 1000 models. The RF ranked the same top four amino acids in the same order as LR (**Figure 6C**).

A further way to identify important features is to examine the sign and amplitude of the coefficients of the features in the linear equations of the LR models. The coefficients in the LR models contain information on the importance of each feature on BA, however, the values cannot be directly used to infer their importance yet. To evaluate the feature importance based on the coefficients in the LR model, we need to standardize each feature by scaling each to have zero mean and standard deviation of one. We also need calculate the variance inflation factor (VIF) for each feature in the models to examine the multicollinearity among the features. The results showed that the VIF values for the coefficients in the minimum AIC model was very low for all the amino acids (< 5.0), indicating that the model did not suffer from multicollinearity and the regression results were reliable (**Table 5**). In fact, the AA-INs had low correlation between each other (**Supplementary Table S9**). The result showed that the coefficients of tyrosine, glycine, serine, and arginine were negative with p < 0.05, suggesting that increasing the INs of these amino acids leads to increased BA. The impact to BA follows the order of tyrosine, glycine, serine, and arginine, which agrees with the results from the correlation between AA-INs with experimental ΔG (**Table 4**).

**Table 5.** Coefficients in the linear equation predicting ΔG for the AA-INs model with minimum AIC before and after feature standardization.

| AA-INs | VIF | Coefficient before standardization | Coefficient after standardization | *p-value* |
| --- | --- | --- | --- | --- |
| Tyr-IN | 1.21 | -0.1535 | -1.0180 | $p=0.000$ |
| Gly-IN | 1.27 | -0.1288 | -0.6530 | $p=0.001$ |
| Ser-IN | 1.36 | -0.0840 | -0.4636 | $p=0.019$ |
| Arg-IN | 1.07 | -0.0805 | -0.5147 | $p=0.004$ |
| Val-IN | 1.09 | -0.0684 | -1.0180 | $p=0.082$ |
| Ile-IN | 1.04 | 0.0731 | 0.3462 | $p=0.045$ |
| Constant | 1.00 | -6.46 | -9.5582 | $p=0.000$ |

### Comparative studies show that the AA-INs models have more consistent performance across datasets than other contact models

As we previously mentioned, the performance of the PRODIGY LR model on the PDBbind dataset has been shown to be worse than its performance on the original PRODIGY data. To examine if the models generated in this work suffered from the same limitation, we split the data used here into three sets: the PRODIGY dataset (Set 1), the PDBbind dataset (Set 2), and the combined dataset (Set 3). We then examined the performance of the minimum AIC models for each of the five feature sets (residue ICs/NIS, atomic ICs/NIS, residue/atomic ICs/NIS, residue/atomic ICs/HS/NIS, and AA-INs) on Set 1 and Set 2 when trained on the combined data set (Set 3). Similar to previous results using the PRODIGY model [25], the R of the correlation between predicted and experimental ΔG decreased greatly on the PDBbind dataset for the residue ICs/NIS model (which is highly similar to the PRODIGY model) and the atomic ICs/NIS (**Table 6**). The combined models, residue/atomic ICs/NIS and residue/atomic ICs/HS/NIS models had a less extreme drop in performance, while the AA-INs had the most similar R across Sets 1 and 2. To supplement these results, we trained each of the 5 groups of models on Set 1, Set 2, and Set 3 individually and calculated correlations between experimental and predicted ΔG for the original sets and after fourfold cross-validation for the minimum AIC models (**Table 7**). These results mirrored those discussed for Table 6, with the AA-INs model having similar performance across the three data sets. Thus, the AA-INs based LR models were more robust to the training dataset than the other contact-based models. Unlike the residue contact models that do not have the identities of the amino acid, the AA-INs model directly includes the amino acids involved in contacts when making predictions.

**Table 6.** Comparison of R values calculated for the PRODIGY dataset (Set 1) and PDBbind dataset (Set 2) using minimum AIC models trained on the combined dataset.

| Model | Number of features used in the model | Set 1 | Set 2 |
| --- | --- | --- | --- |
| Residue ICs/NIS | 5 | 0.69 | 0.54 |
| Atomic ICs/NIS | 2 | 0.61 | 0.49 |
| Residue/ atomic ICs/NIS | 7 | 0.69 | 0.59 |
| Residue/atomic ICs/HS/NIS | 9 | 0.71 | 0.60 |
| AA-INs | 6 | 0.69 | 0.65 |

**Table 7.**
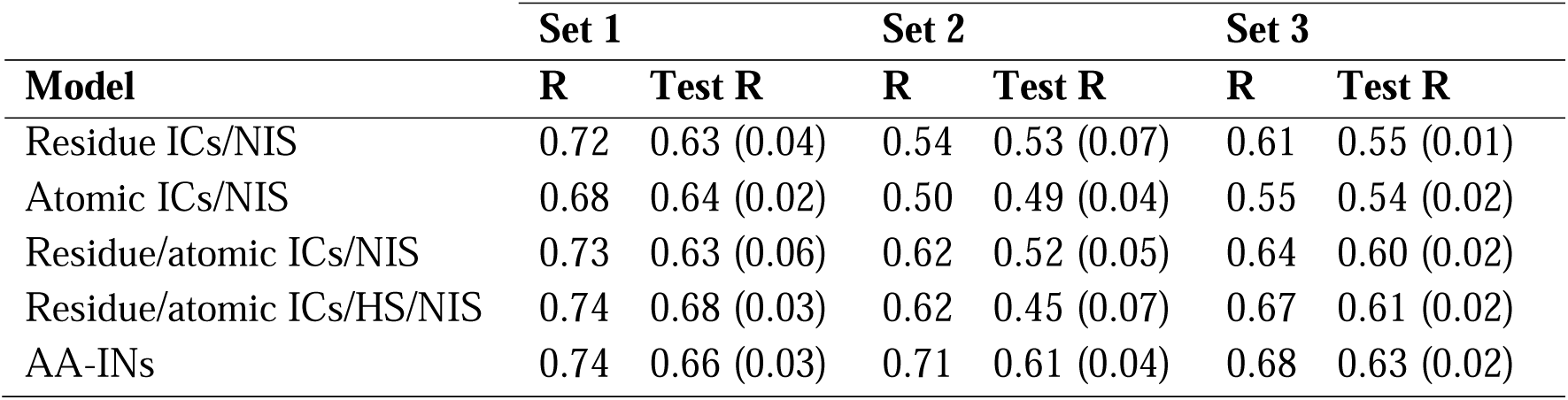
Comparison of the performance of the minimum AIC models using the PRODIGY dataset (Set 1), PDBbind dataset (Set 2), and the combined dataset (Set 3). R is the Pearson’s correlation coefficient of predictions after training on the respective complete datasets. Test R is Pearson’s correlation coefficient (mean with standard deviation) of the test set after 10x fourfold cross-validation.

| Model | Set 1 |  | Set 2 |  | Set 3 |  |
| --- | --- | --- | --- | --- | --- | --- |
|  | R | Test R | R | Test R | R | Test R |
| Residue ICs/NIS | 0.72 | 0.63 (0.04) | 0.54 | 0.53 (0.07) | 0.61 | 0.55 (0.01) |
| Atomic ICs/NIS | 0.68 | 0.64 (0.02) | 0.50 | 0.49 (0.04) | 0.55 | 0.54 (0.02) |
| Residue/atomic ICs/NIS | 0.73 | 0.63 (0.06) | 0.62 | 0.52 (0.05) | 0.64 | 0.60 (0.02) |
| Residue/atomic ICs/HS/NIS | 0.74 | 0.68 (0.03) | 0.62 | 0.45 (0.07) | 0.67 | 0.61 (0.02) |
| AA-INs | 0.74 | 0.66 (0.03) | 0.71 | 0.61 (0.04) | 0.68 | 0.63 (0.02) |

It is worth mentioning that our simple AA-INs model also outperforms many other models. In the previous work, Vangone and Bonvin used 79 complexes from the PRODIGY dataset and compared the performance of the PRODIGY model with 15 previously published models such as DFIRE, CP_PIE and FIREDOCK that were built with different methods (etc. global surface, buried surface area, composite scoring function) [19]. They showed that the PRODIGY model outperformed all these models, suggesting the same for our AA-INs model as it has comparable performance as the PRODIGY model on the PRODIGY dataset. Our AA-INs model also outperforms ISLAND and PPI-Affinity, two recently reported top models, in making predictions of the PRODIGY dataset, as the amino acid sequence-based ISLAND had R = 0.38 and the contact molecular descriptor-based PPI-Affinity had R = 0.62 on the PRODIGY dataset [25].

## Discussion

Despite advancements in understanding PPIs in the past two decades, understanding of how BA is controlled by physical/chemical parameters and how these parameters can be used to make predictions about complex binding affinities remains a challenge. One key set of features that has been commonly used to build models that predict binding affinity from a complex’s structure are the numbers and identities of the protein-protein contacts that hold the complex together. However, inconsistent performance of contact-based models indicates that there is a need for further investigations into the relationship between binding affinity and the contacts in protein-protein complexes. To better understand this relationship, we compiled and studied a dataset of 141 protein-protein complexes with diverse functions and a wide range of BAs and used it to build and test a series of LR models with a range of contact features. These features include both residue-based contact features that have been commonly used in previous BA prediction models and other features (atomic contacts, hydrogen bonding, hydrophobic indices, and the identities of the amino acid at the binding interface) that have received little or no attention in previous models.

In general, we found that the different sets of features provided roughly similar performance in predicting experimental binding affinities. First, a large number of single features produced a Pearson’s correlation R between experimental and predicted BA of around -0.45. The single features that were able to achieve this performance were from all the major contact feature groups investigated here and included total residue ICs and polar/apolar residue ICs; total, polar/apolar, and polar/polar atomic ICs, Tyr-IN, Gly-IN, and Ser-IN. Second, the performance of both minimum AIC and all-feature models for the different investigated feature sets were roughly comparable, with the models for the different feature sets providing R values between 0.55 to 0.65 after fourfold cross-validation. Additionally, continuing to add new features to the models typically resulted in no or very slight improvements in performance. For example, despite the relatively large single feature R values for all the atomics-based contact classes, the minimum AIC atomic ICs model contained only two features (%NIS_polar and polar-apolar ICs). Thus, adding polar-polar and apolar-apolar atomic contacts to this two-feature model was not able to improve the model, despite the high single feature R of atomic polar-polar and apolar-apolar contacts. One possible exception to this trend was that combining atomic and residue ICs resulted in a modest increase in performance, and the future development of models containing both atomic and residue ICs would be of interest. Taken together, these results indicate that LR modeling of BA based on contact features may have an upper limit of an R between predicted and experimental BA of ∼ 0.7. Improving models beyond this limit will likely require the addition of non-contact features or novel ways of describing the contacts at the binding interface.

One of the key results of this study was the determination that the numbers of amino acids in protein-protein contacts (AA-INs) could be used to produce models that have more consistent performance across datasets than models built with standard contact features. Indeed, the AA-INs model had the highest correlation with BA of any of the model groups investigated here, and, in particular, it had the most consistent performance across the two different benchmark datasets (***Table 7***). Using a diverse set of 141 complexes from two benchmark datasets, we generated a best performing AA-INs model with a simple and explicit linear equation involving six amino acids (tyrosine, glycine, serine, arginine, valine, and isoleucine).

We identified four top amino acids that underlie the protein-protein binding strength, which were tyrosine, glycine, serine, and arginine. We showed that tyrosine played the leading role in predicting BA, with the highest correlation of all amino acids between INs and ΔG (R = - 0.49). Its impact surpassed that of the top polarity-classified residue ICs, polar-apolar residues (R = -0.44) and charged-apolar residues (R = -0.39). Using the interfacial number of tyrosine alone, we can generate a LR model to predict BA with R of 0.5. Tyrosine is known to have unique physicochemical properties that makes it the most effective amino acid in mediating molecular recognition [36]. Tyrosine is also the second most enriched amino acid at the binding interface (**Figure 4C** and **Supplementary Table S6**). While tyrosine is a large amino acid, glycine and serine, two of the other top 4 amino acids, are small residues. Thus, these small residues may provide space and flexibility for tyrosine to mediate molecular contacts [36]. Tyrosine and arginine, together with tryptophan, are the three amino acids with the highest enrichment at the interface (1.99× for tyrosine, 1.65× for tryptophan, and 1.61× for arginine) (**Supplementary Table S6**). These three amino acids have been shown to appear in hot spots of the complex interface with a frequency over 10% (21% for tryptophane, 13% for arginine, and 12% for tyrosine) [34]. However, the interfacial number of tryptophan has low correlation with BA. Thus, tryptophan was not found to contribute to interface energetics in our analysis although it is highly enriched at interface and shares many attributes with tyrosine.

In summary, there has been decades of major effort at anatomizing protein-protein interfaces to better understand PPIs and develop therapeutic drugs. The results from this study further our understanding of PPIs by unraveling a quantitative link between the abundance of specific amino acids in contacts at the interface and the strength of PPIs. Thus, interfacial amino acids-based features, such as AA-INs, should be considered in future attempts to build ML models to predict BA for drug design and screening in the pharmaceutical industry or for other applications.

## Additional files

### Supplementary file 1

### Supplementary file 2

This file shows the following information for the 141 complexes in the dataset: PDB IDs, interacting chains, experimental methods to measure Kd, function class, the number of the residue ICs for charged-charged pairs, charged-polar pairs, charged-apolar pairs, polar-polar pairs, polar-apolar pairs, and apolar-apolar pairs, the number of atomic ICs for polar-polar pairs, polar-apolar pairs, and apolar-apolar pairs, the number of hydrogen bonds, hydrophobicity score, % NIS_polar, %NIS_apolar, %NIS_charged, experimental ΔG and the predicted ΔG for the minimal AIC model of the residue ICs/NIS models, atomic ICs/NIS models, residue/atomic ICs/NIS models, and residue/atomic ICs/HS/NIS models.

### Supplementary file 3

This file shows the interfacial numbers of the twenty amino acids for the 141 complexes in the dataset and the predicted ΔG by the minimal AIC AA-INs model.

## Acknowledgements

We thank Kaya Marascalco for the help on compiling the dataset. We also thank Dr. Qianyi Chen for discussions on the RF algorithm and the calculation of the number of hydrogen bonds.

