## Supplemental file 1 for "Predictive Models and Impact of Interfacial Contacts and Amino Acids on Protein-protein Binding Affinity"

A

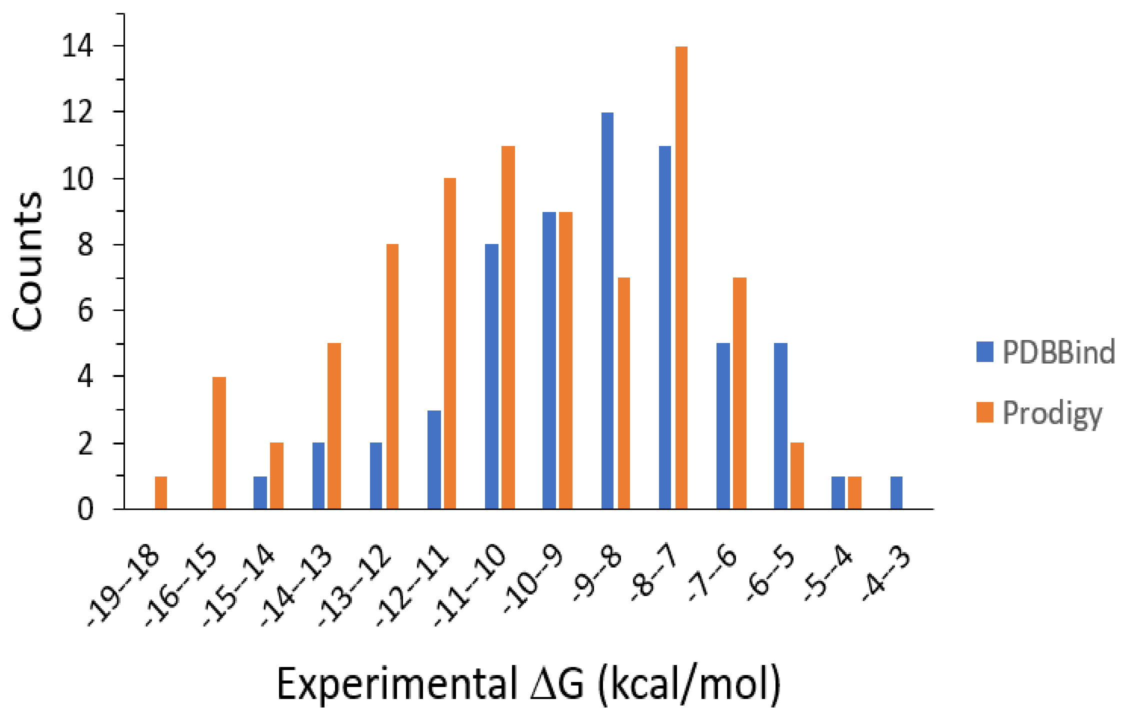


B

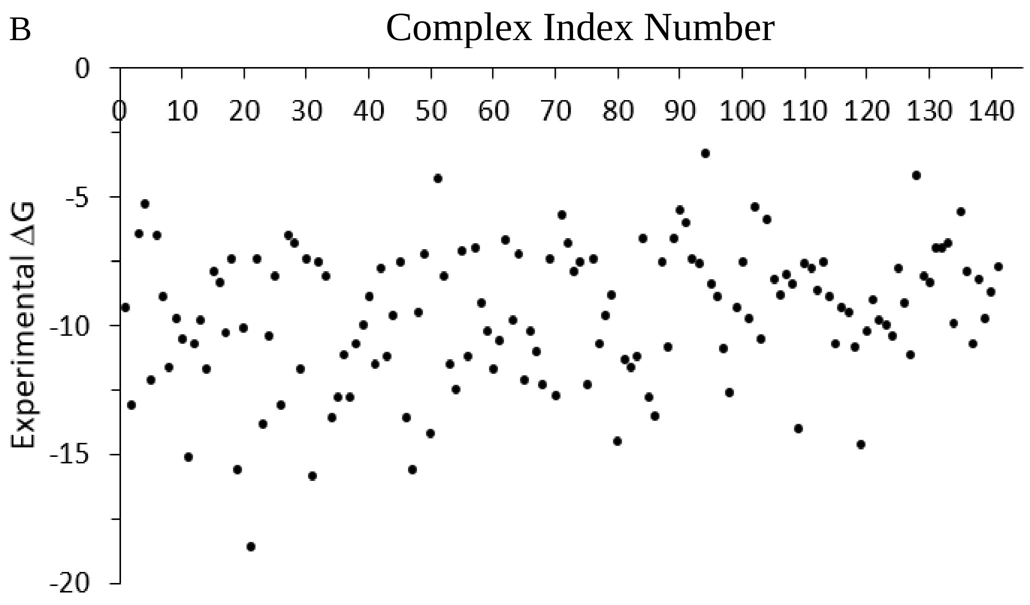


**Figure S1**. (A) Comparision of the histograms of the experimental ΔG for the two datasets. (B) Distribution of the experimental ΔG for the combined dataset of 141 complexes.

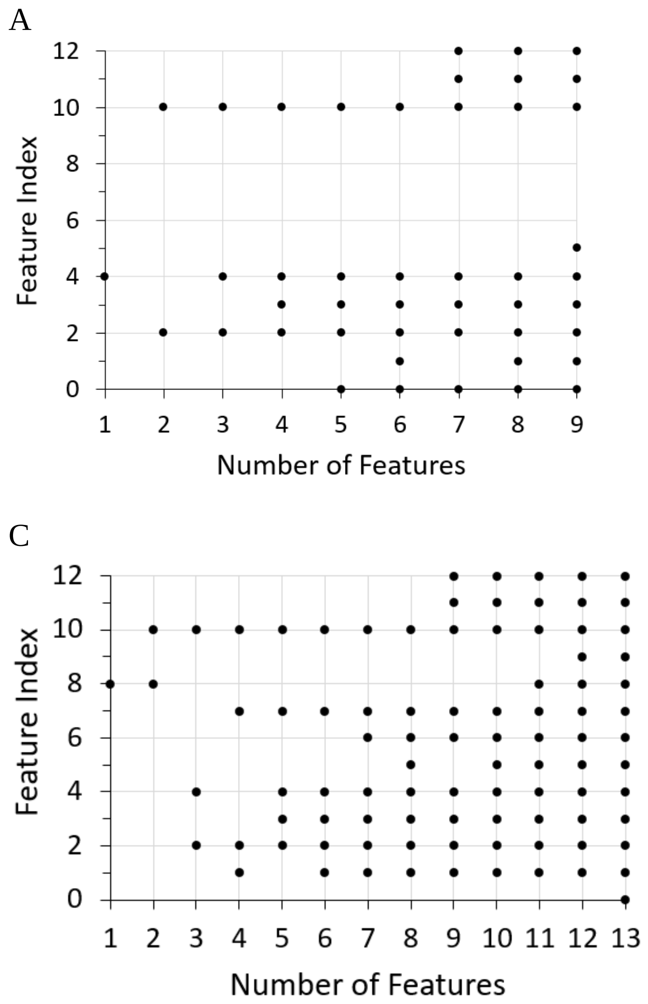


**Figure S2**. Feature compositions in the minimum AIC model for each feature combination. (A) residue ICs/NIS models. (B) Atomic ICs/NIS models. (C) Residue/atomic ICs/NIS models. (D) Residue/atomic ICs/HS/NIS models.

**Table S1.** Pearson's correlation coefficient (R) between experimental ΔG and total number of residue contacts (Res) and atomic contacts (Atom) for the PRODIGY, PDBbind, and their combined datasets.

| **Experimental method** | **Number of complexes in PRODIGY** | **R** | **Number of complexes in PDBbind** | **R** | **Number of complexes in the combined dataset** | **R** |
|---|---|---|---|---|---|---|
| SPR | 39 | -0.59 Res<br>-0.54 Atom | 19 | -0.27 Res<br>-0.24 Atom | 58 | -0.45 Res<br>-0.43 Atom |
| ITC | 20 | -0.54 Res<br>-0.62 Atom | 35 | -0.33 Res<br>-0.38 Atom | 55 | -0.39 Res<br>-0.43 Atom |
| Spectroscopy | 14 | -0.57 Res<br>-0.67 Atom | 8 | -0.34 Res<br>-0.19 Atom | 22 | -0.36 Res<br>-0.32 Atom |
| Stopped-flow | 8 | -0.60 Res<br>-0.65 Atom | 0 | N/A<br>N/A | 8 | -0.60 Res<br>-0.65 Atom |
| Others | 0 | N/A<br>N/A | 28 | -0.008 Res<br>0.06 Atom | 28 | -0.008 Res<br>0.06 Atom |
| Total | 81 | | 90 | | 171 | |

**Table S2**. Composition of the combined dataset used in this study. AB: Antibody/antigen. EI: enzyme/inhibitor. ES: enzyme/substrate. ER: enzyme/regulatory subunit. OR: membrane receptor containing complex. OG: G-protein containing complex. NC: non-cognate complex. OX: miscellaneous complexes.

| | **Classification** | **Number of complexes in PRODIGY set** | **Number of complexes in PDBbind set** | **Number of complexes in the combined set** |
|---|---|---|---|---|
| By function | AB | 12 | 1 | 13 |
| | EI | 12 | 11 | 23 |
| | ES | 7 | 4 | 11 |
| | ER | 4 | 7 | 11 |
| | OR | 8 | 16 | 24 |
| | OG | 11 | 6 | 17 |
| | NC | 8 | 0 | 8 |
| | OX | 19 | 15 | 34 |
| By experimental method | SPR | 39 | 18 | 57 |
| | ITC | 20 | 34 | 54 |
| | spectroscopy | 14 | 8 | 22 |
| | Stopped-flow | 8 | 0 | 8 |
| Total complexes | | 81 | 60 | 141 |

**Table S3**. Comparison of the expected and actual proportion of residue pairs in the contact region of protein-protein complexes.

| | **Expected (%)** | **Actual (%)** |
|---|---|---|
| ICs_charged-charged | 7.6% | 10.0% |
| ICs_charged-polar | 12.0% | 12.1% |
| ICs_charged-apolar | 28.0% | 23.2% |
| ICs_polar-polar | 4.7% | 5.3% |
| ICs_polar-apolar | 22.0% | 20.8% |
| ICs_apolar-apolar | 25.7% | 28.7% |

**Table S4**. Coefficients of features in the linear equations predicting ΔG for of all-feature models.

| | **Residue ICs /NIS model** | | **Atomic ICs /NIS model** | | **Residue/atomic ICs /NIS model** | | **Residue/atomic ICs /HS/NIS model** | |
|---|---|---|---|---|---|---|---|---|
| | Coefficient | *p-value* | Coefficient | *p-value* | Coefficient | *p-value* | Coefficient | *p-value* |
| Residue ICs_charged-charged | -0.0888 | *p=0.110* | - | - | -0.0202 | *p=0.770* | 0.0467 | *p=0.509* |
| Residue ICs_charged-polar | 0.0440 | *p=0.463* | - | - | 0.1236 | *p=0.073* | 0.1697 | *p=0.014* |
| Residue ICs_charged-apolar | -0.1365 | *p=0.002* | - | - | -0.1117 | *p=0.016* | -0.1289 | *p=0.005* |
| Residue ICs_polar-polar | 0.1295 | *p=0.156* | - | - | 0.1972 | *p=0.035* | 0.2880 | *p=0.003* |
| Residue ICs_polar-apolar | -0.1552 | *p=0.001* | - | - | -0.1109 | *p=0.036* | -0.1130 | *p=0.028* |
| Residue ICs_apolar-apolar | 0.0007 | *p=0.983* | - | - | 0.0256 | *p=0.531* | 0.0768 | *p=0.076* |
| HB | - | - | 0.0897 | *p=0.120* | 0.1017 | *p=0.095* | 0.1203 | *p=0.043* |
| Atomic ICs_polar-polar | - | - | -0.0480 | *p=0.105* | -0.0793 | *p=0.014* | -0.0772 | *p=0.014* |
| Atomic ICs_polar-apolar | - | - | -0.0055 | *p=0.634* | 0.0061 | *p=0.595* | 0.0084 | *p=0.454* |
| Atomic ICs_apolar-apolar | - | - | -0.0075 | *p=0.297* | -0.0037 | *p=0.660* | -0.0055 | *p=0.498* |
| % NIS_polar | -0.1522 | *p=0.000* | -0.1397 | *p=0.000* | -0.1466 | *p=0.000* | -0.137 | *p=0.000* |
| % NIS_apolar | -0.0167 | *p=0.560* | -0.0341 | *p=0.216* | -0.0218 | *p=0.441* | -0.0217 | *p=0.428* |
| % NIS_charged | -0.0184 | *p=0.435* | -0.0115 | *p=0.608* | -0.0184 | *p=0.440* | -0.0259 | *p=0.265* |
| HS | - | - | - | - | - | - | -0.1215 | *p=0.003* |
| Constant | -0.0019 | *p=0.000* | -0.0019 | *p=0.000* | -0.0019 | *p=0.000* | -0.0019 | *p=0.000* |

**Table S5**. Pearson's correlation R among the 14 features.

| **Feature No.** | **0** | **1** | **2** | **3** | **4** | **5** | **6** | **7** | **8** | **9** | **10** | **11** | **12** | **13** |
|---|---|---|---|---|---|---|---|---|---|---|---|---|---|---|
| 0 | 1 | 0.50 | 0.41 | -0.03 | 0.01 | -0.12 | 0.67 | 0.63 | 0.56 | 0.36 | -0.35 | -0.02 | 0.37 | 0.48 |
| 1 | | 1 | 0.46 | 0.28 | 0.29 | -0.03 | 0.69 | 0.74 | 0.67 | 0.47 | 0.03 | -0.16 | 0.08 | 0.58 |
| 2 | | | 1 | -0.10 | 0.20 | 0.31 | 0.54 | 0.56 | 0.64 | 0.61 | -0.05 | 0.05 | 0.02 | 0.36 |
| 3 | | | | 1 | 0.57 | -0.06 | 0.23 | 0.36 | 0.28 | 0.15 | 0.41 | -0.21 | -0.27 | 0.43 |
| 4 | | | | | 1 | 0.36 | 0.37 | 0.52 | 0.56 | 0.55 | 0.38 | -0.04 | -0.37 | 0.48 |
| 5 | | | | | | 1 | 0.07 | 0.17 | 0.33 | 0.59 | -0.05 | 0.31 | -0.15 | 0.30 |
| 6 | | | | | | | 1 | 0.90 | 0.84 | 0.61 | 0.04 | -0.08 | 0.02 | 0.66 |
| 7 | | | | | | | | 1 | 0.93 | 0.68 | 0.09 | -0.09 | -0.03 | 0.73 |
| 8 | | | | | | | | | 1 | 0.81 | 0.07 | -0.02 | -0.06 | 0.71 |
| 9 | | | | | | | | | | 1 | -0.04 | 0.09 | -0.03 | 0.57 |
| 10 | | | | | | | | | | | 1 | -0.34 | -0.78 | 0.12 |
| 11 | | | | | | | | | | | | | -0.32 | -0.02 |
| 12 | | | | | | | | | | | | | 1 | -0.10 |
| 13 | | | | | | | | | | | | | | 1 |

**Table S6.** Proportion (%) of each amino acid in the entire complex and at interface as well as the enrichment factor (ratio of the percentage at interface to percentage in entire complex). Data were calculated from all the 141 complexes in the dataset.

| Amino acid | % in entire complex | % at interface | Enrichment factor |
|---|---|---|---|
| Ala | 6.7 | 4.4 | 0.66 |
| Arg | 4.4 | 6.9 | 1.61 |
| Asn | 4.8 | 5.0 | 1.04 |
| Asp | 5.7 | 5.2 | 0.91 |
| Cys | 1.9 | 1.3 | 0.71 |
| Gln | 4.3 | 5.2 | 1.20 |
| Glu | 6.2 | 6.2 | 1.00 |
| Gly | 7.0 | 5.2 | 0.75 |
| His | 2.1 | 3.0 | 1.42 |
| Ile | 5.1 | 4.8 | 0.95 |
| Leu | 8.9 | 8.0 | 0.89 |
| Lys | 6.2 | 6.4 | 1.05 |
| Met | 1.9 | 2.6 | 1.37 |
| Phe | 3.9 | 4.9 | 1.25 |
| Pro | 4.6 | 3.9 | 0.84 |
| Ser | 7.3 | 6.0 | 0.83 |
| Thr | 6.5 | 5.4 | 0.83 |
| Trp | 1.6 | 2.6 | 1.65 |
| Tyr | 3.8 | 7.5 | 1.99 |
| Val | 7.2 | 5.4 | 0.75 |

**Table S7.** Pearson's correlation R between AA enrichment factor (EF) and experimental ΔG.

| **AA-EF** | **R** | **p-value** |
|---|---|---|
| Tyr-EF | -0.30 | <0.0001 |
| Glu-IN | -0.24 | 0.002 |
| Arg-EF | -0.21 | 0.006 |
| Cys-IN | -0.14 | 0.049 |
| Ser-EF | -0.13 | 0.062 |
| Gly-EF | -0.12 | 0.078 |
| Asp-EF | -0.10 | 0.119 |
| Asn-EF | -0.07 | 0.204 |
| Leu-IN | -0.05 | 0.278 |
| Thr-EF | -0.01 | 0.453 |
| Pro-IN | 0.01 | 0.453 |
| Trp-EF | 0.02 | 0.407 |
| Met-IN | 0.05 | 0.278 |
| Val-IN | 0.09 | 0.144 |
| Lys-IN | 0.08 | 0.173 |
| His-IN | 0.10 | 0.119 |
| Phe-IN | 0.14 | 0.049 |
| Ile-IN | 0.18 | 0.016 |
| Gln-IN | 0.24 | 0.002 |
| Ala-IN | 0.28 | 0.0004 |

**Table S8**. Test R for the fourfold cross-validation of the AA-INs model with minimal AIC.

| | **Test R** |
|---|---|
| Trial 1 | 0.66 |
| Trial 2 | 0.62 |
| Trial 3 | 0.62 |
| Trial 4 | 0.65 |
| Trial 5 | 0.60 |
| Trial 6 | 0.65 |
| Trial 7 | 0.64 |
| Trial 8 | 0.64 |
| Trial 9 | 0.63 |
| Trial 10 | 0.62 |
| **Average** | **0.63** |
| Stdev | 0.02 |

**Table S9**. Pearson's correlation R between AA-INs features.

| | **Ala** | **Arg** | **Asn** | **Asp** | **Cys** | **Gln** | **Glu** | **Gly** | **His** | **Ile** | **Leu** | **Lys** | **Met** | **Phe** | **Pro** | **Ser** | **Thr** | **Trp** | **Tyr** | **Val** |
|---|---|---|---|---|---|---|---|---|---|---|---|---|---|---|---|---|---|---|---|---|
| Ala | 1 | 0.08 | 0.00 | 0.05 | 0.17 | 0.00 | 0.08 | 0.05 | -0.05 | -0.02 | 0.06 | 0.11 | 0.17 | 0.09 | 0.21 | 0.09 | 0.10 | -0.10 | -0.10 | 0.11 |
| Arg | | 1 | 0.22 | 0.37 | 0.10 | -0.07 | 0.26 | 0.08 | 0.02 | -0.07 | 0.10 | -0.03 | 0.08 | 0.14 | 0.16 | 0.23 | 0.20 | 0.22 | -0.01 | 0.10 |
| Asn | | | 1 | 0.22 | -0.15 | 0.05 | 0.22 | 0.25 | -0.08 | -0.09 | 0.01 | 0.17 | 0.07 | 0.04 | 0.05 | 0.39 | 0.31 | 0.17 | 0.28 | -0.03 |
| Asp | | | | 1 | -0.07 | 0.09 | 0.21 | 0.12 | -0.24 | 0.05 | -0.05 | 0.32 | 0.04 | 0.10 | 0.12 | 0.20 | 0.09 | 0.15 | 0.15 | -0.01 |
| Cys | | | | | 1 | -0.04 | -0.16 | 0.12 | 0.12 | 0.00 | 0.05 | 0.04 | 0.04 | 0.05 | 0.15 | 0.10 | -0.06 | 0.14 | -0.07 | 0.04 |
| Gln | | | | | | 1 | 0.06 | -0.02 | 0.14 | 0.21 | 0.13 | 0.09 | -0.09 | 0.13 | 0.04 | -0.07 | 0.02 | -0.08 | -0.04 | 0.27 |
| Glu | | | | | | | 1 | -0.06 | 0.03 | -0.08 | 0.05 | 0.47 | 0.20 | 0.16 | 0.18 | 0.20 | 0.19 | -0.03 | 0.12 | 0.12 |
| Gly | | | | | | | | 1 | 0.11 | 0.03 | 0.02 | -0.14 | 0.03 | -0.11 | 0.02 | 0.42 | 0.26 | 0.21 | 0.29 | 0.04 |
| His | | | | | | | | | 1 | 0.01 | -0.05 | -0.03 | 0.16 | 0.13 | 0.10 | 0.09 | -0.03 | 0.11 | 0.02 | 0.10 |
| Ile | | | | | | | | | | 1 | 0.16 | 0.09 | 0.05 | 0.20 | -0.11 | 0.02 | -0.05 | -0.11 | 0.02 | 0.18 |
| Leu | | | | | | | | | | | 1 | 0.05 | 0.11 | 0.06 | 0.00 | 0.14 | 0.08 | -0.17 | -0.17 | 0.21 |
| Lys | | | | | | | | | | | | 1 | 0.18 | 0.17 | 0.04 | 0.08 | -0.06 | -0.12 | 0.04 | 0.02 |
| Met | | | | | | | | | | | | | 1 | 0.10 | -0.02 | 0.08 | -0.08 | 0.01 | -0.17 | 0.05 |
| Phe | | | | | | | | | | | | | | 1 | 0.17 | 0.12 | 0.01 | -0.09 | -0.14 | 0.29 |
| Pro | | | | | | | | | | | | | | | 1 | 0.10 | 0.12 | 0.09 | -0.03 | 0.11 |
| Ser | | | | | | | | | | | | | | | | 1 | 0.30 | 0.21 | 0.25 | 0.20 |
| Thr | | | | | | | | | | | | | | | | | 1 | -0.02 | 0.26 | 0.14 |
| Trp | | | | | | | | | | | | | | | | | | 1 | 0.31 | -0.13 |
| Tyr | | | | | | | | | | | | | | | | | | | 1 | -0.02 |
| Val | | | | | | | | | | | | | | | | | | | | 1 |
